# Single-cell metabolic dynamics of early activated CD8 T cells during the primary immune response to infection

**DOI:** 10.1101/2020.01.21.911545

**Authors:** Lauren S. Levine, Kamir J. Hiam, Diana M. Marquez, Iliana Tenvooren, Diana C. Contreras, Jeffrey C. Rathmell, Matthew H. Spitzer

## Abstract

Memory T cells conventionally rely on oxidative phosphorylation and short-lived effector T cells on glycolysis. Here, we investigate how T cells arrive at these states during an immune response. In order to understand the metabolic state of rare, early activated T cells, we adapted mass cytometry to quantify metabolic regulators at single-cell resolution in parallel with cell signaling, proliferation, and effector function. We interrogated CD8 T cell activation in vitro as well as the trajectory of CD8 T cells responding to *Listeria monocytogenes* infection, a well-characterized *in vivo* model for studies of T cell differentiation. This approach revealed a unique metabolic state in early activated T cells characterized by maximal expression of glycolytic and oxidative metabolic proteins. Peak utilization of both pathways was confirmed by extracellular flux analysis. Cells in this transient state were most abundant five days post-infection before rapidly downregulating metabolic protein expression. This approach should be useful for mechanistic investigations of metabolic regulation of immune responses.

## Introduction

Understanding the regulatory mechanisms underlying immune responses is crucial to developing more rationally designed treatment strategies for acute and chronic infections, autoimmune diseases and malignancy (Buck et al., 2017). CD8 T cells, when activated, expand and differentiate into potent short-lived effector cells (SLECs) as well as long-term memory cells, which confer durable protection against re-infection and cancer relapse (Badovinac et al., 2007; Callahan et al., 2016; Restifo et al., 2012). The former mediate primary adaptive immune responses against pathogens through the release of cytotoxic granules and pro-inflammatory cytokines (Araki et al., 2010; Pearce et al., 2009). In contrast, long-lived memory cells remain quiescent until re-encountering antigen, upon which they rapidly mediate secondary immune responses (Gerriets and Rathmell, 2012). The field of immunometabolism has provided critical insight into these processes, revealing a complex regulatory interplay of signaling, metabolic and epigenetic adaptations during CD8 T cell differentiation (Olenchock et al., 2017; Zhang and Romero, 2017).

Upon activation, effector CD8 T cells undergo clonal expansion, necessitating as many as 20 replication cycles to generate sufficient daughter cells to clear pathogens (Badovinac et al., 2007). This process is energetically costly and requires rapid ATP production for the biosynthesis of essential building blocks (Zhang and Romero, 2017). Previous studies suggest that the exit from quiescence is supported by a dramatic metabolic shift from oxidative phosphorylation (OXPHOS) in naïve cells, fueled by beta-oxidation of long chain fatty acids (LCFA), to aerobic glycolysis in SLECs, characterized by lactate production in the setting of adequate oxygen (Calderon et al., 2018; Menk et al., 2018; Wang et al., 2011). This metabolic conversion permits continued cycling through the pentose phosphate pathway and thus generation of intermediates necessary for nucleic acid and lipid biosynthesis. This adaptation also circumvents negative feedback induced by the accumulation of pyruvate and acetyl-CoA (Lee et al., 2014; Zhang and Romero, 2017). Additional feed-forward mechanisms supporting this process include the activation of transcription factors downstream of phosphoinositide 3-kinase (PI3K) signaling. For instance, hypoxia inducible factor 1 (HIF1α) mediates the upregulation of nutrient receptors including glucose transporter 1 (Glut1), the main point of entry for glucose into T cells (Wang et al., 2011).

Meanwhile, the transition to the memory T cell fate is associated with the inhibition of PI3K/mTORC1 signaling and silencing of aerobic glycolysis by AMP-activated protein kinase (AMPK). Instead, AMPK favors mitochondrial biogenesis and fusion (Borges da Silva et al., 2018; Buck et al., 2016; D’Souza et al., 2007; Pearce et al., 2009), which is mediated by peroxisome proliferator-activated receptor gamma coactivator 1-alpha (PGC1α) (Andrejeva and Rathmell, 2017; Calderon et al., 2018). This tightly regulated metabolic shift results in an LCFA-fueled oxidative program characterized by increased mitochondrial mass (Buck et al., 2016). This property of memory cells confers additional oxidative potential, known as spare respiratory capacity (SRC), to permit more rapid recall during secondary immune responses (van der Windt et al., 2012).

While many studies of bulk T cell populations suggest that a reciprocal, tightly regulated relationship exists between OXPHOS and glycolysis and the signaling cascades that regulate these pathways, their precise interactions in individual cells have yet to be elucidated. Moreover, the regulation of metabolic machinery in rare, early activated T cells remains poorly understood. The early stages of infection lead to antigen specific CD8 T cells acquiring transient cell states preceding differentiation into effector subsets, but precisely how these intermediate stages of differentiation metabolically orchestrate rapid proliferation and differentiation has remained technically challenging (Joshi et al., 2007; Kalia et al., 2010; Obar and Lefrançois, 2010). Recently, considerable advances in single-cell analysis have enabled studies of signaling and effector programs in T cells at high resolution (Krishnaswamy et al., 2014; Mingueneau et al., 2014). Analogous studies of T cell metabolic regulation would likely provide new insights. For instance, a recent study utilizing stable isotope tracing in activated T cells has found that OXPHOS may be more prominent in effector T cells *in vivo* than was previously thought (Ma et al., 2019). However, in the absence of single-cell resolution, it remains unclear whether the same cells are responsible for both OXPHOS and glycolysis, or alternatively, whether individual cells already differentiate and preferentially utilize one pathway versus the other during the effector phase. Many of the regulatory mechanisms that govern cellular metabolism are post-transcriptional and therefore not directly measurable by RNA-sequencing (Andrejeva and Rathmell, 2017). Therefore, a single-cell proteomic approach provides unique opportunities.

Mass cytometry utilizes metal-tagged antibodies to directly measure up to 45 proteins simultaneously in individual cells (Bandura et al., 2009; Bendall et al., 2011). This approach has permitted characterization of various aspects of cellular behavior including phenotype, signaling (Bodenmiller et al., 2012), proliferation (Good et al., 2019) and chromatin state (Cheung et al., 2018). Here, we have further adapted this platform to measure expression levels of enzymes and transporters involved in metabolic checkpoints. We have integrated direct quantitative evaluation of the signaling cues thought to mediate their regulation along with proteins indicative of CD8 T cell fate and function. In this study we utilize this approach to interrogate key inflection points of the CD8 T cell response to *Listeria monocytogenes* infection (Lm-OVA), a well-characterized model of CD8 T cell differentiation (McGregor et al., 1970).

## Results

### Mass cytometry permits high-dimensional quantification of metabolic regulators in single CD8 T cells

T cell differentiation requires the coordinated interplay of signaling and metabolic pathways, including the upregulation of rate-limiting enzymes and regulatory switches. The transition to aerobic glycolysis in SLECs is mediated by co-stimulatory signaling through CD28 via the AKT/PI3K pathway (Pollizzi et al., 2015; Wang et al., 2011); therefore, we measured the downstream intermediates mTOR, pS6, p4EBP1, and HIF1α (Fig. 1A, Table S1). Signaling through this pathway promotes glucose uptake through the Glut1 receptor and the transcription of glycolytic enzymes (Dennis et al., 2012), including glyceraldehyde-3-phosphate dehydrogenase (GADPH) (Fig. 1B), a critical metabolic switch implicated in glycolytic activity, which we also quantified.

**Figure 1.**
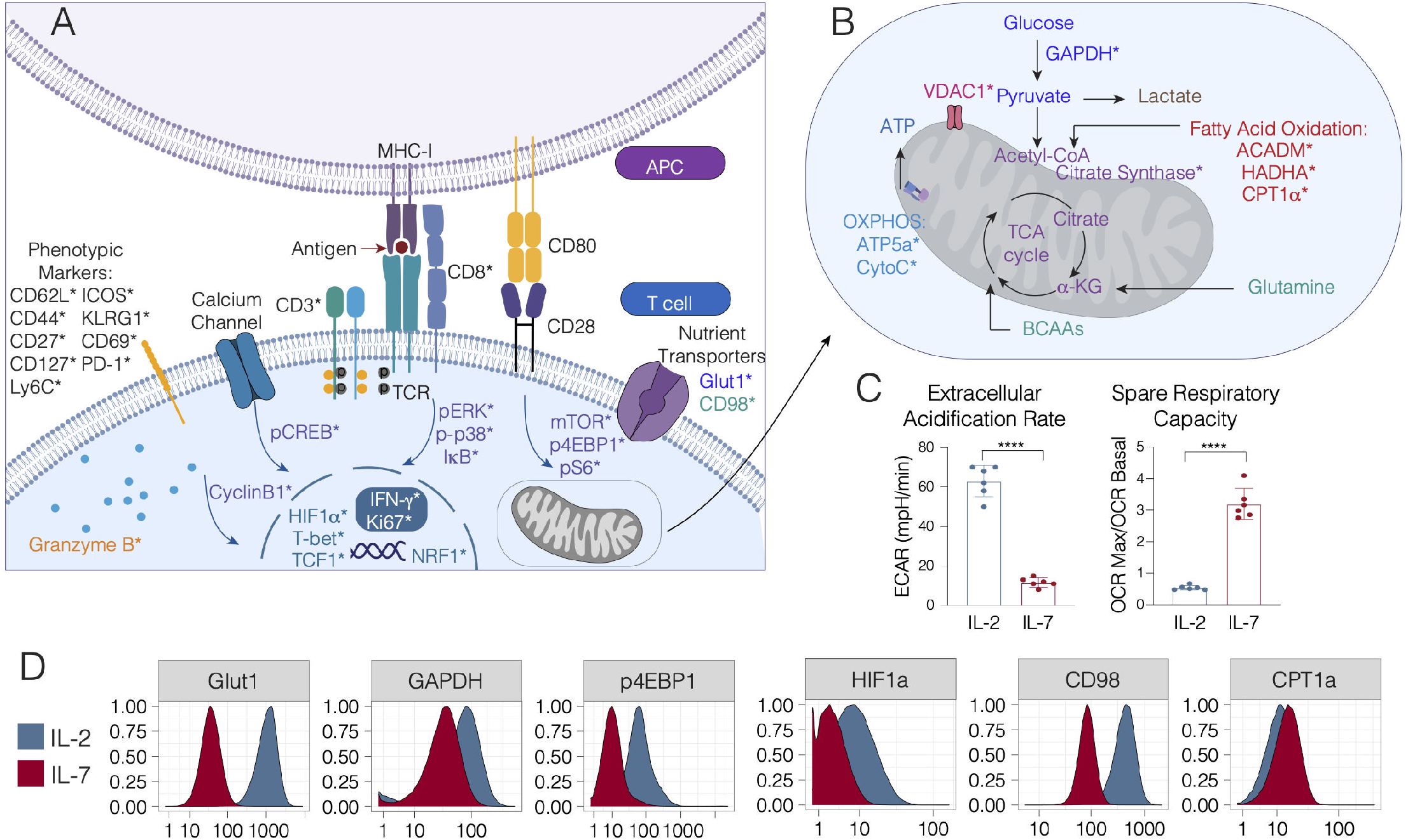
Querying the integrated functional program of CD8 T cell activation. Panel schematic depicting signaling, metabolic, effector and phenotypic targets interrogated by mass cytometry. Cell surface, cytosolic and nuclear markers are depicted in **(A)** and mitochondrial markers are denoted in **(B)**. Markers directly measured by mass cytometry are demarcated by an asterisk (*). OT-1 transgenic CD8 T cells were stimulated with cognate peptide (SIINFEKL) in the presence of IL-2 (100U/ml) for 72 hours, followed by 3 washes to remove antigen and polarization in IL-2 or IL-7 (both 10ng/mL) to generate OT-1_eff_ or OT-1_mem_. Samples were harvested at Day 7 for metabolic analysis by mass cytometry and Seahorse assay analysis by Mitochondrial Stress Test. **(C)** Mass cytometry expression profiles of Day 7 OT1_eff_ and OT1_mem_ for key metabolic enzymes as depicted by histograms. **(D)** Extracellular acidification rate and Oxygen consumption rate by Seahorse Assay depicted in bar plots and with significance analysis by student’s t-test (p<.001 ****). Error bars represent SEM. Data are representative of 3 independent experiments.

To investigate how the TCA cycle is regulated in activated T cells, we evaluated the expression of citrate synthase (CS) (Fig. 1B, Table S1), the first step of the cycle, which is directly regulated by NAD+/NADH ratio, ADP/ATP ratio and succinyl-coA levels (Wiegand and Remington, 1986). As branched chain amino acid metabolism has been demonstrated to be critical for effective T cell activation (Ren et al., 2017), we sought to understand this process by measuring the large neutral amino acid transporter (LAT1) chaperone CD98 (Fig. 1A, Table S1), a key mediator of the import of these essential nutrients (Hayashi et al., 2013; Nii et al., 2001; Sinclair et al., 2013).

Previous work has described a reciprocal relationship between aerobic glycolysis and OXPHOS, the latter of which is associated with memory T cell differentiation. Therefore, we sought to understand this regulation at the single-cell level by measuring CPT1α, an enzyme that catalyzes the transport of LCFA from the cytoplasm to the mitochondria and that is critical for memory T cell function (van der Windt et al., 2012). Additionally, we measured the mitochondrial trifunctional complex, also known as hydroxyacyl-CoA dehydrogenase (HADHA), which catalyzes the final three steps of LCFA oxidation to acetyl-CoA in the mitochondria (Carpenter et al., 1992). As the role of β-oxidation of medium-chain fatty acids in T cell function has not been extensively evaluated (Howie et al., 2018), we also measured the expression of medium-chain acyl-CoA dehydrogenase (ACADM), an essential enzyme that catalyzes the initial step of this process (Fig. 1B, Table S1). Moreover, we measured key components of the electron transport chain, including cytochrome C (CytoC) and ATP synthase (ATP5a) (Fig. 1B, Table S1). To understand the counterregulatory processes governing OXPHOS activity and overall energy, we measured voltage-dependent ion channel 1 (VDAC1), a critical regulator controlling cytoplasmic-mitochondrial cross-talk (Fig. 1B, Table S1)(Cunningham et al., 2018; Tarze et al., 2007).

The cell signaling pathways that mediate mitochondrial fusion and biogenesis include MAP kinase and NFkB, which are activated during T cell priming (Calderon et al., 2018; Enamorado et al., 2018; Laforge et al., 2016); therefore, we measured the levels of phosphorylated (p) ERK and p-p38 MAP kinases in addition to the total levels of NFkB inhibitor alpha (IkBa). Calcium signaling, triggered by TCR ligation, has also been implicated in this process (Feske, 2007; Fracchia et al., 2013). Therefore, we additionally measured pCREB levels (Fig. 1A, Table S1).

It has been proposed that the activity of metabolic pathways induces the activity epigenetic regulators such as Ezh2, which directly impact T cell fate and function (Chisolm et al., 2017; Gray et al., 2017). Therefore, we included a full range of well-characterized surface markers and transcription factors to subset T cells into naïve, central memory, effector memory and terminal effector populations. Finally, to measure the impact on all of these factors on T cell proliferation during clonal expansion, we measured expression of cyclinB1 and Ki67. To assess production of cytotoxic mediators, we also measured granzyme B (Fig. 1A, Table S1).

### Mass cytometry recapitulates metabolic phenotypes of CD8 T cell differentiation in vitro

In order to query the metabolic program underlying antigen-specific CD8 T cell activation *in vitro,* we first stimulated TCR transgenic OT-1 splenocytes in the presence of their cognate antigen (the SIINFEKL peptide from ovalbumin) and IL-2 for 72 hours. After this initial priming period, antigen was removed, and cells were polarized in IL-2 or IL-7 for an additional 4 days to generate effector (OT-1_eff_) or central memory cells (OT-1_mem_), as described previously (Carrio et al., 2014; Pearce et al., 2009; van der Windt et al., 2012). We analyzed the resulting cells by mass cytometry and real-time metabolic profiling by Seahorse assay (Fig. 1C, S1B-D). In keeping with prior studies (Pearce et al., 2009; van der Windt et al., 2012), OT-1_eff_ exhibited higher rates of extracellular acidification associated with glycolytic activity, while OT-1_mem_ possessed marked spare respiratory capacity (Fig. 1C).

Consistent with these results, OT-1_eff_ expressed elevated levels of glycolytic proteins at day 7 of activation, as evidenced by robust upregulation of Glut1 and GAPDH (Fig. 1D, Fig S1A), suggestive of active glucose uptake and utilization. The expression of targets of the PI3K/mTORC1 pathway, including p4EBP1 and HIF1α, were likewise elevated in OT1_eff_ (Fig. 1D), consistent with the promotion of aerobic glycolysis. Also in keeping with previous data (Ren et al., 2017), the amino acid transporter CD98 was more highly expressed in OT1_eff_ relative to OT-1_mem_ (Fig. 1D). In contrast to their effector counterparts, OT-1_mem_ did not demonstrate this glycolytic profile, but instead upregulated CPT1 α (Fig. 1D, S1A), which promotes OXPHOS in memory T cells (van der Windt et al., 2012).

### *Dynamic metabolic changes in canonical subsets of activated CD8 T cells* in vivo

To understand the metabolic changes during CD8 T cell differentiation in a more physiologic context, we next evaluated the trajectory of the response to acute infection *in vivo.* C57BL/6 mice were infected with *Listeria monocytogenes* expressing whole cytoplasmic ovalbumin (Lm-OVA), a well-characterized model of CD8 T cell differentiation and metabolism (Buck et al., 2016; Pearce et al., 2009; van der Windt et al., 2012). Splenocytes were harvested daily over the first nine days post-infection for analysis by mass cytometry. We began by identifying canonical T cell differentiation states and investigating changes in metabolic enzyme and transporter expression over the course of the immune response (Fig. 2A, S2A). Unsupervised clustering analysis revealed considerable heterogeneity and dynamic functional changes across all major canonical T cell subsets over the course of the primary immune response to *Listeria monocytogenes* (Fig. 2A-B).

**Figure 2.**
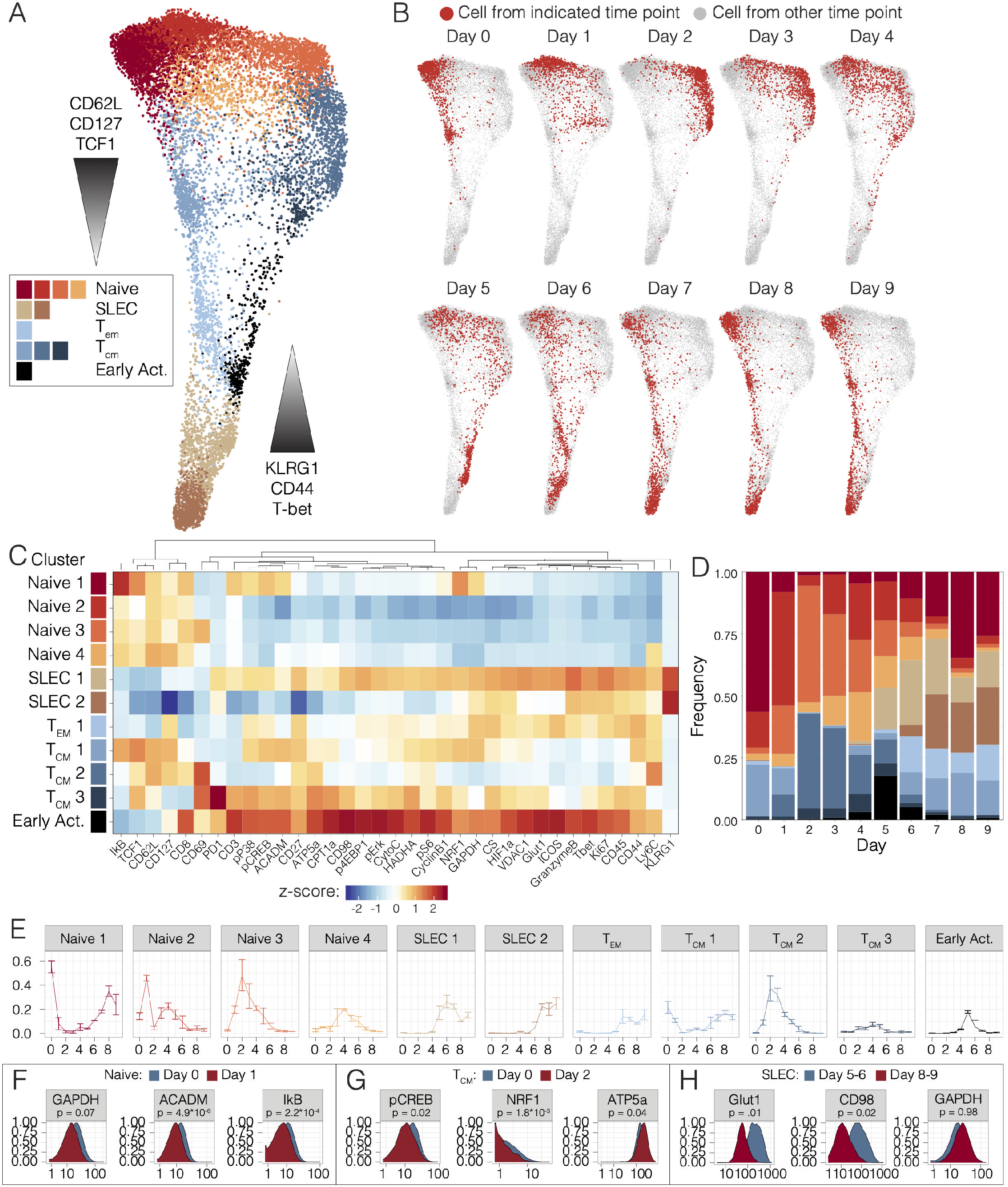
Single-cell analysis of the CD8 T cell effector program *in vivo.* **(A)** Pooled CD8 T cells from mice at days 0 to 9 of Lm-OVA infection (n=2-3 mice per time point) clustered by Phenograph and visualized by a force-directed graph. **(B)** Force-directed graphs indicating cellular distribution by time point. **(C)** Functional and phenotypic median expression profiles for each CD8 T cell cluster. **(D)** Cluster proportion by time point **(E)** Individual cluster frequency profiles at days 0 to 9 p.i. **(F)** Expression profiles of metabolic and signaling in naïve clusters between days 0 and 1 p.i. **(G)** Histograms depicting the expression of functional markers in central memory cells between days 0 and 2 p.i. **(H)** Metabolic expression profiles of SLEC clusters between days 5-6 and 8-9 p.i. Significance analysis of all medians by two-tailed student’s t-test (p<0.05 *, p<0.01 **) is displayed.

At baseline, most naïve cells were predominantly contained within cluster Naïve 1, characterized by the expression of ACADM, pCREB, p-p38, NRF1 and weak expression of GAPDH (Fig. 2C). However, three new clusters, Naïve 2 and Naïve 3, emerged at days 1 and 2 post infection (p.i) (Fig. 2D-E), all characterized by the downregulation of all of the above metabolic and signaling markers (Fig. 2C, S2A). Interestingly, these clusters demonstrated low IkB expression, suggestive of signaling through NFkB pathway (Fig. 2C, 2F, S2A-B). While most naïve T cells were contained within the Naïve 2 cluster at day 1 post-infection (p.i.) (Fig. 2D-E), this gave way to a predominance of the Naïve 3 cluster and days 2 and 3 p.i. (Fig. 2D-E).

By day 4 p.i., all these new clusters as well as an additional cluster, Naïve 4, were present in similar proportions (Fig. 2D-E). Notably, the Naïve 1 cluster began to re-emerge at day 6 p.i., and ultimately dominated the naïve pools from day 7 p.i. onwards (Fig. 2D-E). This predominance was associated with the involution of clusters Naïve 2, Naïve 3, and Naïve 4, which became nearly undetectable by day 7 p.i (Fig. 2E). These findings are consistent with activation of both bystander and antigen specific T cells in the early stages of acute infection (Chu et al., 2013; Jiang et al., 2003) but reveal the metabolic adaptations that these cells undertake. Overall, these data support previous reports of a metabolically quiescent profile of naïve T cells, but they suggest heterogeneity and transitions within even these cells.

Evaluation of the central memory cells over the course of infection revealed a similar pattern, starting with cluster T_CM_1, characterized by intermediate expression of expected markers of LCFA and OXPHOS including p-p38, pCREB, ACADM, HADHA, NRF1, and dim expression of ATP5a, CPT1 a, pErk and CytoC (Fig. 2C). Interestingly, this cluster also expressed GAPDH and pS6 but dimly expressed HIF1α compared to effector subsets (Fig 2C, S2A). However, days 1-2 p.i., were marked by emergence of cluster T_CM_2 (Fig. 2D-E), which downregulated all of these metabolic and signaling factors, with only weak expression of HADHA and pS6 and upregulation of ATP5a (Fig. 2C, 2G, S2C). This cluster predominated at days 2 through 4 p.i. Notably, day 2 p.i. was also marked the emergence of cluster T_CM_3 (Fig. 2D-E), defined by expression of enzymes of fatty acid oxidation (FAO), including CPT1α, HADHA, ACADM, along with oxidative proteins, such as CS, ATP5a, VDAC1, CytoC (Fig. 2C, S2A). These cells also expressed less pS6 and GAPDH, suggestive of a state predominantly fueled by FAO (Fig. 2C). Commensurate with this oxidative profile, cells in this cluster also expressed p-p38, pErk, and pCREB (Fig. 2C, S2A). While cells in T_CM_3 also demonstrated expression of downstream intermediates of the PI3K cascade, such as p4EBP1 and pS6, along with transcription factors associated with aerobic glycolysis, such as HIF1α, these were associated with lower GADPH expression (Fig. 2C, S2A). This metabolically active T_CM_ cluster was transient, completely regressing by day 7 p.i. (Fig. 2D-E). Notably T_CM_1 reemerged at day 5 p.i. and remained the predominant T_CM_ cluster from days 6 through 9 p.i (Fig. 2D). These data confirm the previously oxidative profile of central memory cells but also reveal dynamic metabolic changes within these subsets over the course of an immune response.

Effector memory cells (T_EM_) uniformly constituted cluster T_EM_1, which emerged at day 5 p.i. and maintained stable frequency through day 9 p.i (Fig. 2D-E). These cells demonstrated a more glycolytic metabolic profile, with upregulation of GAPDH, Glut1 and HIF1a, and dim oxidative and FAO marker expression (Fig. 2C, S2A). Meanwhile, SLECs comprised clusters SLEC1 and SLEC2 and emerged at days 5 and 6 p.i., respectively (Fig. 2D-E). These two clusters demonstrated distinctive metabolic phenotypes. The first population to appear, SLEC1, demonstrated expression of p4EBP1, pS6, HIF1α, Glut1, and GAPDH suggestive of a glycolytic profile (Fig. 2C, 2H, S2A, S2D). Recent studies have demonstrated that early effector cells continue TCA cycle engagement fueled by the uptake of amino acids and LCFA (O’Sullivan et al., 2014; Ren et al., 2017); consistently, cells in this cluster also expressed HADHA, CD98, CS, VDAC1 (Fig. 2C, 2H, S2A, S2D). However, ATP5a and CPT1α levels in this cluster were lower than those observed in the more active T_CM_ clusters, such as T_CM_3, distinguishing them from these more classically oxidative pools (Fig. 2C, S2A). In comparison, cluster SLEC2 demonstrated a more muted metabolic profile, downregulating expression of all metabolic mediators except HIF1α, GAPDH and CS, taking on the terminal glycolytic state observed in previous studies (Fig. 2C, 2H, S2A, S2D). As expected, the more metabolically active cells in cluster SLEC1 expressed higher levels of Ki67 and granzyme B compared to cluster SLEC2 (Fig. 2C, S2A). Taken together, these findings agree with previous reports of a predominantly glycolytic terminal effector state.

### Early activated T cells exhibit maximal expression of glycolytic and oxidative proteins

In addition to these well-characterized cell subsets, unsupervised high-dimensional analysis also revealed a group of early activated T cells that emerged at day 4 post-infection (Fig. 2D-E). These cells had high expression of Ki67, indicative of proliferation, and expressed high levels of CD44, CD27 and ICOS but low levels of CD62L (Fig. 2C, S2A). This early activated cluster was most abundant at day 5, when it comprised nearly 20% of the CD8 T cell population, and it nearly completely disappeared by day 7 (Fig. 2D-E). As ICOS has been found to signal through the PI3K cascade (Zeng et al., 2016), we anticipated that this population would be glycolytic. Indeed, these early activated cells expressed the highest levels of Glut1 and GAPDH across all CD8 T cells (Fig. 2C). However, these cells simultaneously exhibited peak expression of oxidative markers, including CPT1α, HADHA, ACADM, and ATP5a (Fig. 2C, S2A). Commensurate with this observation, the signaling program of this population was marked by maximal expression of both pS6 and pCREB as well as minimal expression of IkB, reflecting simultaneous activity of both the PI3K/mTORC1 and NFkB pathways (Fig. 2C, S2A). In contrast to SLEC and memory cells, these early activated cells also expressed maximal expression of the amino acid transporter CD98 (Fig. 2C, S2A).

Given the unique metabolic expression profile of these early activated cells, we sought to confirm these observations through direct inspection of the primary data. We undertook further phenotypic analysis of these cells, which identified the high-affinity IL-2 receptor subunit, CD25, as another surface marker co-expressed by this population of interest, consistent with their recent activation (Fig. 3A). Consistent with the clustering analysis, we found that these manually gated cells peaked in frequency at day 5 followed by a rapid decline in abundance (Fig. 3B). Moreover, in comparison to all other CD8 T cells present in the animals at day 5, these cells clearly expressed elevated levels of both glycolytic and oxidative proteins (Fig. 3C, S3A). Therefore, early activated T cells exhibited peak expression of metabolic mediators of oxidative and glycolytic pathways.

**Figure 3.**
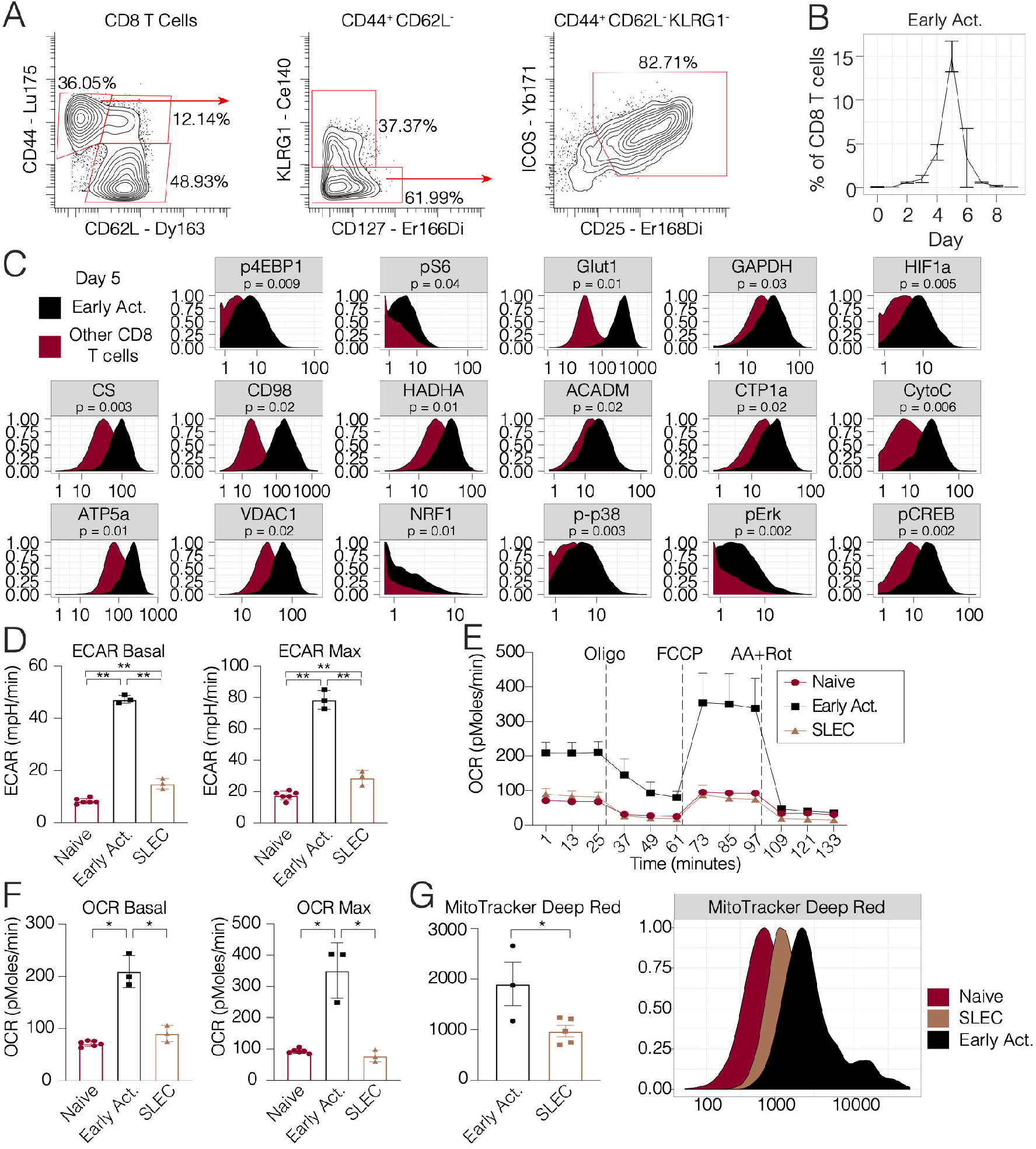
Early activated T cells exhibit a distinctive metabolic profile characterized by peak oxidative and glycolytic activity. **(A)** Biaxial scatter plots indicating surface marker expression profile of the early activated T cell pool. **(B)** Frequency of early activated cells during days 0 to 9 p.i. **(C)** Metabolic expression profiles of metabolic and signaling markers in early activated T cells in comparison to all other CD8 T cells during days 0 to 9 p.i. as depicted by histograms. Significance analysis of all medians by paired two-tailed student’s t-test (p<0.05 *, p<0.01 **) is displayed. CD8 T cell subsets of interest were sorted at days 5 (including, naïve cells, early activated cells) and 8 (SLECs) p.i. and analyzed by Mitochondrial Stress Test (n=5 mice per time point). **(D)** Basal ECAR and maximal ECAR measured upon oligomycin administration**. (E)** OCR over time. **(F)** Basal and maximal OCR readings obtained upon FCCP administration. **(G)** MitoTracker signal in each subset (n=5 mice per subset). Significance analysis by paired twotailed student’s t-test (p<0.05 *, p<0.01 **). Error bars represent SEM. Data are representative of at least 2 independent experiments.

### Early activated T cells demonstrate peak glycolytic activity and increased mitochondrial activity and mass

Since these early activated T cells were distinguished by simultaneously elevated levels of glycolytic and oxidative enzymes, we posited that this expression profile would translate to greater metabolic activity along these pathways when compared to their SLEC counterparts. To assess real-time bioenergetic flux through oxidative and glycolytic pathways, we sorted naïve, early activated, and SLEC T cells for analysis by Seahorse assay (Fig S3B). As expected, SLECs demonstrated significantly higher baseline and maximum ECAR compared to their naïve counterparts (Fig. 3D), confirming a predominantly glycolytic program driving the terminal effector state *in vivo.* However, in accordance with their enzymatic expression profile by mass cytometry, early activated T cells exhibited significantly higher basal and maximal ECAR even compared to SLECs (Fig. 3D).

Moreover, baseline and maximal OCR did not significantly differ between the SLEC and naive pools, as described previously (van der Windt et al., 2012) (Fig. 3E-F). However, early activated T cells did indeed exhibit significantly higher oxidative activity compared to both the SLEC and naïve cells (Fig. 3E-F). Since these cells exhibited maximal expression of CPT1α and electron transport complexes, we hypothesized that that this population would possess SRC, in keeping with previous reports studying memory T cells that upregulate these enzymes (van der Windt et al., 2012). Indeed, while neither the naïve or SLEC pools were capable of surpassing their baseline OCR upon FCCP administration, the OCR of early activated T cells nearly doubled (Fig. 3E-F). As SRC has been associated with greater mitochondrial mass (Buck et al., 2016), we sought to quantify the mitochondrial content of these cells using MitoTracker Deep Red, a fluorescent dye staining mitochondria in live cells. Consistent with our mass cytometry and Seahorse data, the early activated T cells contained significantly more mitochondrial mass than the SLEC or naïve pools based on MitoTracker staining (Fig. 3G). Overall, these observations confirmed the unique, simultaneously oxidative and glycolytic profile in early activated T cells.

### Antigen-specific CD8 T cells transit through the early activation state commensurate with the onset of proliferation

In order to query the antigen-specificity of these metabolic adaptations of early T cell activation, we adoptively transferred OT-1 T cells into congenic hosts, which were then infected with Lm-OVA. Splenocytes were analyzed daily from days 3 through 7 p.i for metabolic analysis by mass cytometry (Fig. 4A-B, S4A). Indeed, unsupervised clustering analysis of adoptively transferred OT-1 T cells revealed early activated cells with an analogous state of metabolic activity, arising in small numbers at day 3 p.i. and peaking at day 4 before rapidly regressing by day 5 (Fig. 4A-B, S4A). The kinetics of the emergence of this cluster were slightly earlier compared to the previously characterized endogenous cells, perhaps a result of higher TCR affinity or increased frequency of antigen-specific precursor cells. Consistent with our findings in endogenous CD8 T cells (Fig. 2C, 3C), cells comprising this cluster exhibited simultaneous peak expression of markers of glycolysis, OXPHOS, and LCFA oxidation (Fig. 4C, S4A). We hypothesized that these metabolic adaptations were undertaken in support of clonal expansion of antigen-specific populations. Therefore, we assessed the proliferation of CFSE-labeled adoptively transferred OT-1 T cells on days 3 through 7 p.i. by flow cytometry. Commensurate with the emergence of this early activated metabolic state, the first antigen-specific T cells to divide did so at day 4 p.i. (Fig. 4D). By day 5 p.i., all adoptively transferred cells had divided multiple times (Fig. 4D). Consistent with this finding, the total number of OT1 T cells only modestly increased between days 3 and 4 p.i., but they subsequently rapidly expanded between days 4 and 5 p.i before plateauing thereafter (Fig. 4E). These observations collectively suggest that early activated antigen-specific CD8 T cells undergo a transition to a metabolic state characterized by peak OXPHOS and glycolytic activity at the same time as they begin blasting, supporting the dramatic expansion of these cells during productive immune responses.

**Figure 4.**
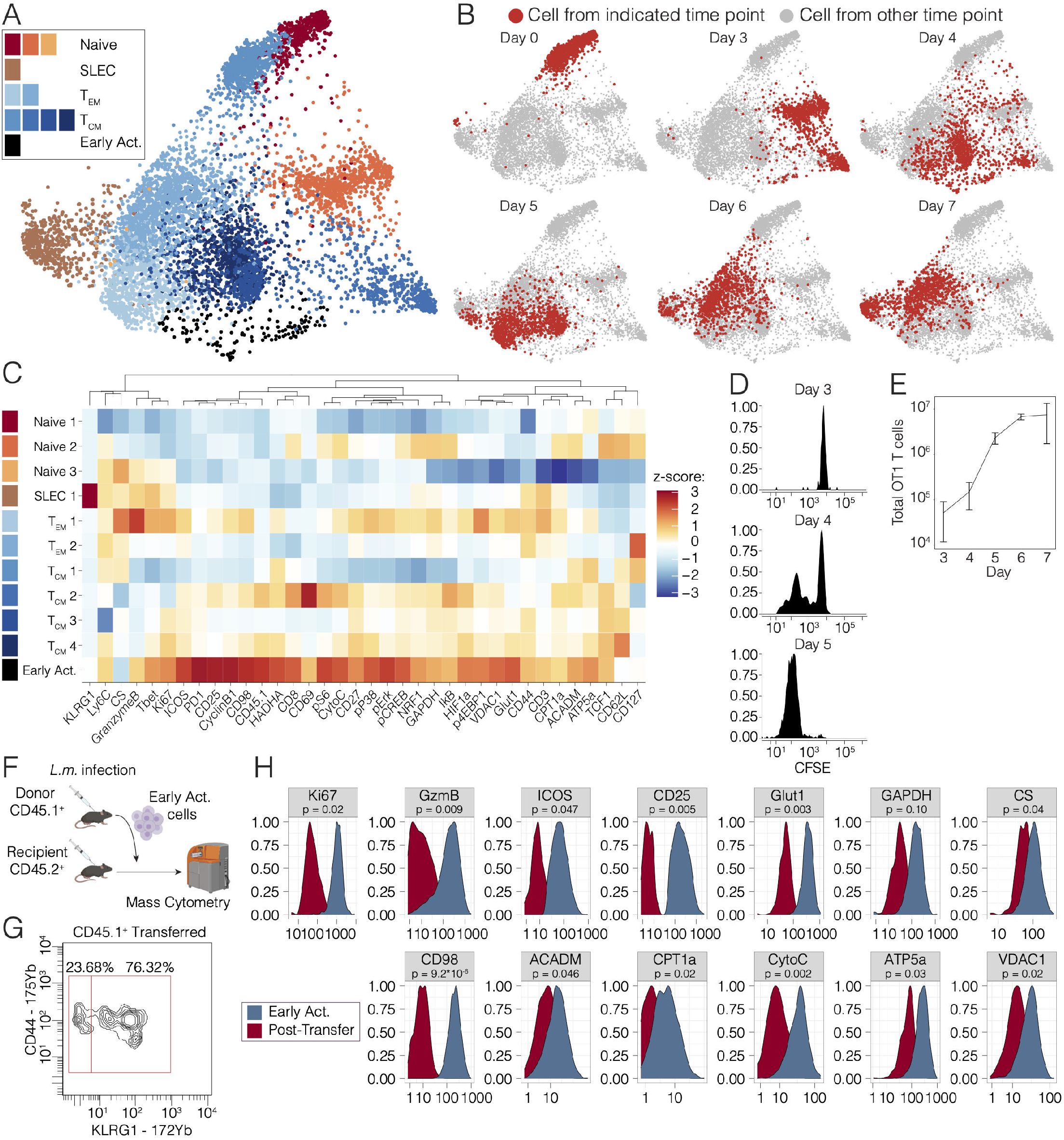
The early activated metabolic state is antigen-specific and transient. OT-1 T cells were adoptively into congenic hosts, which were then infected with 5*10^4^ CFU Lm-OVA. Splenocytes were harvested daily on days 3 through 7 p.i for metabolic analysis by mass cytometry. **(A)** Pooled OT1 cells from mice at days 3 to 7 of Lm-OVA infection (n=1-3 mice per time point) clustered by Phenograph and visualized by a force-directed graph. **(B)** Force-directed graphs indicating cellular distribution by time point. **(C)** Functional and phenotypic median expression profiles for each CD8 T cell cluster**. (D)** Proliferation of adoptively transferred OT1s as measured by CFSE dilution at days 3-5 p.i. and **(E)** absolute cell counts at days 3-7 p.i. **(F)** Early activated T cells were sorted from CD45.1^+^ mice at Day 5 p.i. and transferred into infected CD45.2^+^ hosts at Day 5 p.i (n=2 per group) and sacrificed 4 days later for analysis by mass cytometry **(G)** Differentiation state of the transitional subset determined by CD44 and KLRG1 expression at day 9 p.i. **(H)** Metabolic and signaling marker profiles before and after transfer at days 5 and 9 p.i. are represented by histograms. Significance analysis of the medians by twotailed student’s t-test (p<0.05 *, p<0.01 **) is displayed.

### Transient expression of metabolic proteins in early activated T cells

As early activated CD8 T cells with peak metabolic protein expression were highly transient, only detectable for a few days during the immune response, we investigated the changes that take place in these cells thereafter. We sorted early activated T cells and transferred them into congenic hosts before isolating splenic T cells four days later (Fig. 4F). At the end of this period of time, the transferred early activated cells had given rise to a mixture of cells with phenotypes consistent with SLECs (CD44^+^ KLRG1^+^) as well as memory cells (CD44^+^ KLRG1^-^). Given the transient, elevated expression of cell cycle markers by the early activated cells (Fig. 2C), we hypothesized that these cells would proliferate briefly upon adoptive transfer. Indeed, these cells expanded but lost expression of Ki67 over the course of the four days (Fig. 4H, S4B). These early activated T cells also downregulated expression of CD25 and ICOS as well as Granzyme B by this later time point (Fig. 4H, S4C). Consistent with a transient burst of metabolic activity, these cells exhibited markedly lower expression of both glycolytic and oxidative markers compared to the early activated T cells from which they originated (Fig. 4H, S4C). Collectively, these data indicate that during CD8 T cell differentiation, early activated antigen-specific T cells undergo a transient period of peak metabolic activity. Thereafter, downregulation of glycolytic and oxidative pathways takes place coordinate with differentiation into short-lived or memory cells.

## Discussion

Mass cytometry permits broad-spectrum characterization of immune responses in healthy and diseased states (Spitzer and Nolan, 2016). To date, this approach has been used to query the phenotypic and signaling adaptations undertaken by cells during differentiation (Bendall et al., 2014; Zunder et al., 2015). However, until now, the coordinated downstream metabolic cues supporting these programs have remained incompletely understood at the single-cell level. Here, we directly measured the expression levels of essential nutrient receptors, enzymes, signaling intermediates, and markers of cellular differentiation and effector function at the proteomic level. This allowed us to more thoroughly characterize CD8 T cell responses during acute infection, highlighting the metabolic adaptations of canonical T cell subsets and capturing a unique metabolic state in rare, early activated T cells.

Previous models of the naïve-to-effector transition based on bulk data have proposed a process in which OXPHOS is repressed to promote aerobic glycolysis (Sukumar et al., 2013; Wang et al., 2011). However, a recent study of intracellular flux in activated T cells has reported that effector T cells may utilize oxidative phosphorylation *in vivo* (Ma et al., 2019). Additionally, it has been observed that effector T cells engage in more active LCFA uptake than their memory cell counterparts, which instead have been shown to mobilize these substrates from lysosomal triglycerides (Sullivan et al., 2014). It is therefore feasible that fatty acid uptake may provide additional substrate for OXPHOS early in the course of T cell activation. Our data unify these observations, supporting a coordinated program in which glycolysis and OXPHOS are maintained simultaneously in individual cells during an earlier stage of T cell activation.

Our approach to metabolic profiling by mass cytometry affords investigators the opportunity to functionally characterize the metabolic adaptations of rare cellular populations, such as antigenspecific T cells. These cells would be otherwise difficult to analyze by current standard metabolomics assays due to the prohibitively large number of cells and extensive processing and ex vivo culture techniques required for these studies (Cantor et al., 2017; van der Windt et al., 2016). Here, we were able to characterize the metabolic, signaling and phenotypic progeny of adoptively transferred cells with single-cell resolution. This approach revealed a diversification during CD8 T cell differentiation in the context of acute infection, with a highly metabolically active and proliferative state in T cells early in the course of their response, which later give rise to cells with both memory and terminal effector phenotypes.

The maximal expression of metabolic proteins early after T cell activation suggests a potential role for TCR ligation and/or co-stimulation during CD8 T cell priming. Notably 4-1BB ligation during co-stimulation has been shown to induce mitochondrial fusion via TRAF2-mediated signaling through p38 and PGC1α (Calderon et al., 2018; Enamorado et al., 2018). Similarly, CD28 ligation has been demonstrated to induce CPT1α expression *in vitro* (Klein Geltink et al., 2017). Whether these signals potentiate the observed spare respiratory capacity and increased mitochondrial mass in early activated T cells will be important to determine. As Drp1-mediated mitochondrial fission has been described in effector cells during metabolic reprogramming to the aerobic glycolytic program (Buck et al., 2016), it is possible that the absence of co-stimulation and loss of IL-2 signaling upon pathogen clearance may result in mitophagy and/or mitochondrial fission, repressing oxidative activity in terminal effector subsets.

As the importance of metabolism to immune cell fate and function is increasingly appreciated, methods to evaluate these pathways in models of productive and dysregulated immune responses will be critical. The approach presented here may be adapted to any cell type of interest, including both immune cells and non-immune cells, such as interacting epithelial tissues or tumors. This methodology should enable investigators to query the functional programs underlying the development of the full spectrum of immune cell lineages and their compromised state in the context of autoimmunity or malignancy. Additionally, integrated functional analysis of rare cellular subsets will permit simultaneous evaluation of the effects of various treatments on rare populations, such as tumor infiltrating lymphocytes or neoantigen-specific T cells.

## Acknowledgements

We would like to thank the members of the Spitzer lab, Vinh Ngyuen, Stanley Tamaki, Sagar Bapat, Adil Daud, Lawrence Fong, Andrei Goga, Rushika Perera for their experimental contributions and helpful discussions. Listeria monocytogenes strain 10403s expressing OVA (Lm-OVA) was kindly provided by Shomyseh Sanjabi (UCSF). We acknowledge the Parnassus Flow Cytomtery Core Facility supported in part by NIH Grants P30DK063720, S10OD018040, S10OD018040 and S10OD021822. Recombinant human IL-2 (IL-2; TECIN; Teceleukin) was provided by the National Cancer Institute.

This work was supported by NIH grants DP5OD023056 to M.H.S. and R01DK105550 and R01HL136664 to J.C.R. L.S.L. was supported by Conquer Cancer Foundation Young Investigator Award grant CA-0122026. M.H.S. is a Chan Zuckerberg Biohub investigator and a Parker Institute for Cancer Immunotherapy investigator.

## Author Contributions

L. S.L., K.J.H, J.C.R., and M.H.S. conceptualized this study and designed experiments. L.S.L, K.J.H. D.M.M., I.T. and D.C.C. performed experiments. L.S.L, K.J.H., and M.H.S. performed data analysis. L.S.L. and M.H.S. wrote the manuscript. M.H.S. supervised the study.

## Methods

### Animals

All mice were housed in an American Association for the Accreditation of Laboratory Animal Care-accredited animal facility and maintained in specific pathogen-free conditions. Animal experiments were approved and conducted in accordance with AN157618. Wild-type female C57BL/6 mice and BoyJ CD45.1 between 8-10 weeks old were purchased from The Jackson Laboratory and housed at our facility. TCR Transgenic OT-I CD45.1 mice and heterozygous CD45.2/CD45.1 mice were bred at our facility. Animals were housed under standard SPF conditions with typical light/dark cycles and standard chow.

### Infectious Agents

Listeria monocytogenes strain 10403s expressing OVA (Lm-OVA) was kindly provided by Shomyseh Sanjabi (UCSF). Lm-OVA stocks frozen at −80 C were grown overnight at 37°C in BHI broth supplemented with 5 ug/ml Erythromycin (Bio Basic, Amherst, New York). Then, overnight cultures were sub-cultured by diluting into fresh BHI broth supplemented with 5 ug/ml Erythromycin and grown for 4 hours. Bacteria CFU was then quantified by measuring optical density at 600 nm. Bacteria were then diluted to 5×10^4^ CFU / 100μl in sterile PBS and 100 μl was injected per mouse i.v. via the retroorbital vein.

### Mass Cytometry Antibodies

All mass cytometry antibodies and concentrations used for analysis can be found in Table S1. Primary conjugates of mass cytometry antibodies were prepared using the MaxPAR antibody conjugation kit (Fluidigm, South San Francisco, CA) according to the manufacturer’s recommended protocol sourcing metals from Fluidigm (Fluidigm, South San Francisco, CA) or Trace Sciences International (Richmond Hill, Canada). Following labeling, antibodies were diluted in Candor PBS Antibody Stabilization solution (Candor Bioscience GmbH, Wangen, Germany) supplemented with 0.02% NaN3 to between 0.1 and 0.3 mg/mL and stored long-term at 4°C. Each antibody clone and lot was titrated to optimal staining concentrations using primary murine samples with all appropriate positive and negative controls: polyclonal murine CD8 T cells purified by positive selection kit (Stem Cell Technologies, Vancouver, Canada) stimulated with PMA/Ionomycin via eBioscience Cell Stimulation Cocktail (ThermoFisher Scientific, Waltham, Massachusetts) for 15 minutes, 3 hours and 6 hours or plate-bound CD3 (145-2C11) and soluble CD28 (37.51) antibodies (UCSF Monoclonal Antibody Core, San Francisco) for 3 days, OT-1 splenocytes at day 7 of IL-2 or IL-7 polarization as below, and appropriate CD8 T cell subsets (Naïve, Short-lived Effector and Central Memory) at Day 8 of Lm-OVA infection. Titration results were cross-referenced to the literature as described in the text.

### In vitro OT1 Stimulation and Polarization

OT-1 polarizations were carried out as previously described (Carrio et al, 2004). Briefly, splenocytes from OT-1 mice were cultured at 1*10^6^ cells/mL in 24 well-plates of complete RPMI (UCSF Media Core facility) supplemented with 10% FBS (Omega Scientific, Tarzana, California), 100U/mL penicillin-streptomycin (Fisher Scientific, Hampton, New Hampshire), 2mM L-glutamine (Sigma-Aldrich, St. Louis, Missouri) and 50μM β-mecaptoethanol (Thermo Fisher Scientific, Waltham, Massachusetts) and 10mM HEPES (UCSF Media Core Facility) in the presence of OVA_257-264_ peptide (0.1nM) (Invivogen, San Diego, California) and IL-2 (100U/ml) (Teceleukin) kindly provided by NCI Frederick. After 3 days in culture, activated cells were washed 3 times with RPMI 1640 and recultured in T25 culture flasks at 1*10^5^ cells/mL in the presence of either IL-7, IL-15 (Biolegend, San Diego, California) or IL-2 (Teceleukin) kindly provided by NCI Frederick. (all cytokines 10ng/ml) After 2 additional days in culture, cells were passaged and recultured under the same conditions without peptide for an additional two days for total of 7 days in culture. Viability was confirmed by trypan blue exclusion (Thermo Fisher, Waltham, Massachusetts) or mass cytometry as described below.

### Cell Preparation

All tissue preparations were performed simultaneously from each individual mouse, as previously reported (Spitzer et al. 2017). After euthanasia by CO2 inhalation, spleens were collected and homogenized in PBS + 5mM EDTA at 4°C. All tissues were washed with PBS/EDTA and re-suspended 1:1 with PBS/EDTA and 100mM Cisplatin (Enzo Life Sciences, Farmingdale, NY) for 60s before quenching 1:1 with PBS/EDTA + 0.5% BSA to determine viability as previously described (Spitzer et al., 2015). Cells were centrifuged at 500 g for 5 min at 4°C and re-suspended in PBS/EDTA/BSA at a density between 1-10*10^6^ cells/ml. Care was taken to maintain all samples at 4°C during all phases of tissue harvest and preparation except viability staining and fixation. Suspensions were fixed for 10 min at RT using 1.6% PFA (Fisher Scientific, Hampton, New Hampshire) and frozen at −80°C.

For experiments with adoptively transferred OT1 T cells, immunomagnetic enrichment was performed to facilitate the detection of extremely rare cells before proliferation. Following lysis of red blood cells with ACK lysis buffer (ThermoFisher Scientific, Waltham, Massachusetts), EasySep Streptavidin Negative Selection was used with the following biotinylated antibodies: MHCII (AF6-120.1), CD11c (N418), Ly6C (RB6-8C5), B220 (RA3-6B2), CD4 (GK1.5), Ter119 (TER-119).

### Mass-Tag Cellular Barcoding

Mass-tag cellular barcoding was performed as previously described (Zunder et al., 2015). Briefly, 1*10^6^ cells from each animal were barcoded with distinct combinations of stable Pd isotopes in 0.02% saponin in PBS. Samples from any given tissue from each mouse per experiment group were barcoded together. Cells were washed once with cell staining media (PBS with 0.5% BSA and 0.02% NaN3), and once with 1X PBS, and pooled into a single FACS tube (BD Biosciences, San Jose, California). After data collection, each condition was deconvoluted using a single-cell debarcoding algorithm (Zunder et al., 2015).

### Mass Cytometry Staining and Measurement

Cells were resuspended in cell staining media (PBS with 0.5% BSA and 0.02% NaN3), and antibodies against CD16/32 (BioLegend, San Diego, California) were added at 20 mg/ml for 5 min at RT on a shaker to block Fc receptors. Surface marker antibodies were then added, yielding 500 uL final reaction volumes and stained for 30 min at RT on a shaker. Following staining, cells were washed 2 times with cell staining media, then permeabilized with methanol for at 10 min at 4°C. Cells were then washed twice in cell staining media to remove remaining methanol, and stained with intracellular antibodies in 500 uL for 1 hour at RT on a shaker. Cells were washed twice in cell staining media and then stained with 1mL of 1:4000 191/193Ir DNA intercalator (Fluidigm, South San Francisco, CA) diluted in PBS with 4% PFA overnight. Cells were then washed once with cell staining media, once with 1X PBS and once with Cell Acquisition Solution (Fluidigm, South San Francisco, CA). Care was taken to assure buffers preceding analysis were not contaminated with metals in the mass range above 100 Da. Mass cytometry samples were diluted in Cell Acquisition Solution containing bead standards (see below) to approximately 10^6^ cells per mL and then analyzed on a Helios mass cytometer (Fluidigm, South San Francisco, CA) equilibrated with Cell Acquisition Solution. We analyzed 1-5*10^5^ cells per animal per time point, consistent with generally accepted practices in the field. For adoptive transfer experiments, 1-4*10^6^ cells per animal were analyzed.

### Mass Cytometry Bead Standard Data Normalization

Data normalization was performed as previously described (Spitzer et al., 2017). Briefly, just before analysis, the stained and intercalated cell pellet was resuspended in freshly prepared Cell Acquisition Solution containing the bead standard at a concentration ranging between 1 and 2*10^4^ beads/ml. The mixture of beads and cells were filtered through a filter cap FACS tubes (BD Biosciences) before analysis. All mass cytometry files were normalized together using the mass cytometry data normalization algorithm (Finck et al., 2013), which uses the intensity values of a sliding window of these bead standards to correct for instrument fluctuations over time and between samples.

### Adoptive T Cell Transfer

For adoptive transfer of transitional cells and SLECs, T cells were sorted by FACS from splenocytes harvested from WT CD45.2 C47Bl/6 mice or CD45.1 BoyJ mice 5 days postinfection. Then, viable sorted cells were counted by hemocytometer and Trypan blue staining, resuspended in sterile PBS and transferred into infection-matched congenic mice intravenously via the retroorbital vein.

For adoptive transfer of pathogen specific T cells to validate the antigen specificity of transitional cells, CD8 T cells were immunomagnetically enriched from the spleens of CD45.1 OT1 TCR transgenic mice with EasySep Streptavidin Negative Selection using the following biotinylated antibodies: MHCII (AF6-120.1), CD11c (N418), Gr1 (RB6-8C5), B220 (RA3-6B2), CD4 (GK1.5), Ter119 (TER-119). Viable cells were quantified by counting on a hemocytometer with Trypan blue staining. 1×10^6^ Cells were then resuspended in sterile PBS and transferred into naïve WT CD45.2 mice intravenously via the retroorbital vein.

### Flow Cytometry and Cell Sorting

Cells were stained for viability with Zombie-NIR dye. Cell surface staining was performed in cell staining media (PBS with 0.5% BSA and 0.02% NaN3) for 30 minutes at room temperature. The following anti-mouse antibodies were used: TCRβ – APC (H57-597), CD8 – PE (53-5.8), CD62L – BV421 (MEL-14), KLRG1 – BV510 (2F1/KLRG1), CD44 – PE-Cy7 (IM7), CD25 – FITC (3C7), CD19 – APC-Cy7 (1D3/CD19), F480 APC-Cy7 (BM8). Stained cells were analyzed with an LSR II flow cytometer (BD Biosciences). MitoTracker Deep Red (Thermo Fisher, Waltham, Massachusetts) staining was performed per manufacturer’s instructions and as previously (Scharping et al, 2016). For MitoTracker Deep Red experiments, Zombie-UV dye was used (Biolegend, San Diego, California).

For sorting experiments, cells were prepared as described for flow cytometry and then sorted into FBS containing media (RPMI 1640, 20% FBS, 1% HEPES, 100 mg/mL penicillin/streptomycin) on a FACSAria II (BD Biosciences).

### Seahorse Assays

Seahorse Assays were carried out utilizing Agilent Mitochondrial Stress Test kit as previously (van der Windt et al, 2012) and per the manufacturer’s instructions. Oxygen consumption rates (OCR) and extracellular acidification rates (ECAR) were measured in XF media (non-buffered RPMI 1640 containing 10 mM glucose, 2mM L-glutamine, and 1 mM sodium pyruvate) under basal conditions and in response to 1 μM oligomycin, 1 μM fluoro-carbonyl cyanide phenylhydrazone (FCCP) and 100 nM rotenone + 1 μM antimycin A (all from Agilent, Santa Clara, California) using a 96 well XF Extracellular Flux Analyzer (EFA) (Agilent, Santa Clara, California).

### Statistical Analysis

All significance analysis of Seahorse data and cellular frequency was performed by paired twosided student’s t-test with error bars representing SEM in Prism v8. (GraphPad, San Diego, California). Analysis of median protein expression was performed by paired or unpaired (as indicated) two-sided student’s t-test in R.

### Unsupervised Clustering Analysis

Cell clusters were identified using the Phenograph algorithm as implemented in the ‘cytofkit’ package in R. Standard settings were utilized (with k = 30 for endogenous CD8 T cells and k = 100 for OT1 T cells).

### Data Visualization

Unsupervised force-directed graphs were generated as previously reported (Spitzer et al., 2015) with the following modifications. Single cells were downsampled to n = 1,000 cells from each condition. All cells were combined in a single graph with edge weights defined as the cosine similarity between the vectors of marker values of each cell. All the pairwise distances were calculated and for each node only the 10 edges of highest weight were retained. The graph was then laid out using the ForceAtlas2 algorithm in Gephi.

### Data Availability

Mass cytometry data will be made publicly available as a report on Cytobank (www.cytobank.org) with linked flow cytometry standard (.fcs) files upon acceptance of the manuscript.

---

**Figure S1.**
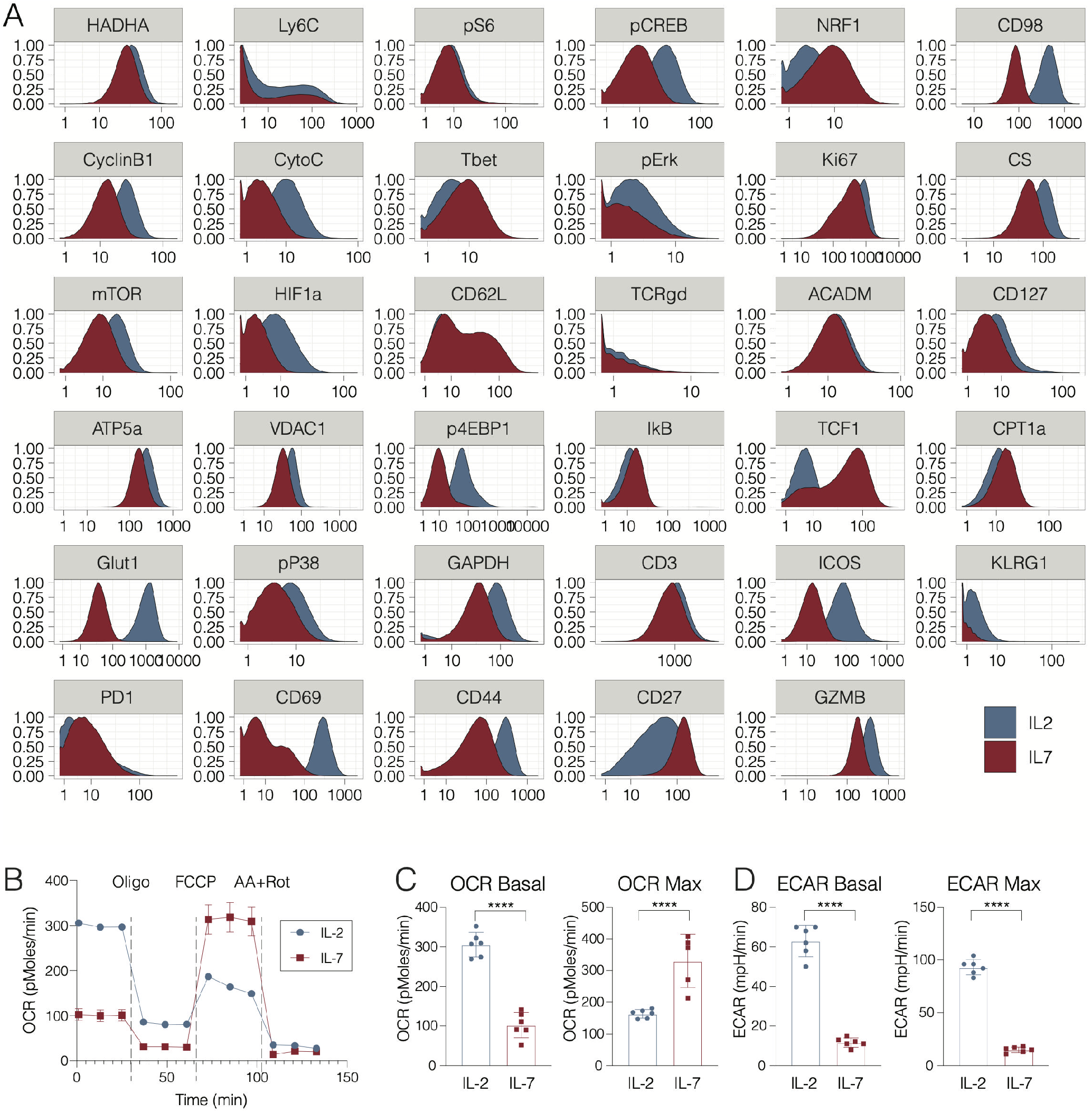
Assessing the integrated functional program of antigen specific CD8 T cell activation in vitro. (A) OT-1 transgenic CD8 T cells were stimulated with cognate peptide (SIINFEKL) in the presence of IL-2 (100U/ml) for 72 hours, followed by 3 washes to remove antigen and polarization in IL-2, IL-7 or IL-15 (all 10ng/mL) to generate OT-1_eff_ or OT-1_mem_. Samples were fixed for mass cytometry at all time points depicted and cells were harvested at Day 7 for Seahorse assay analysis by Mitochondrial Stress Test. Mass cytometry expression data for key metabolic, signaling and effector markers of interest. (B) OCR tracings as quantified by Seahorse. (C) Basal and maximal OCR readings and (D) basal and maximal ECAR readings as quanfied by Seahorse. Significance analysis by paired two-tailed student’s t-test (p<0.05 *, p<0.01 **, p=0.0001***, p<0.0001****). Error bars represent SEM. Data are representative of at least 2 independent experiments.

**Figure S2.**
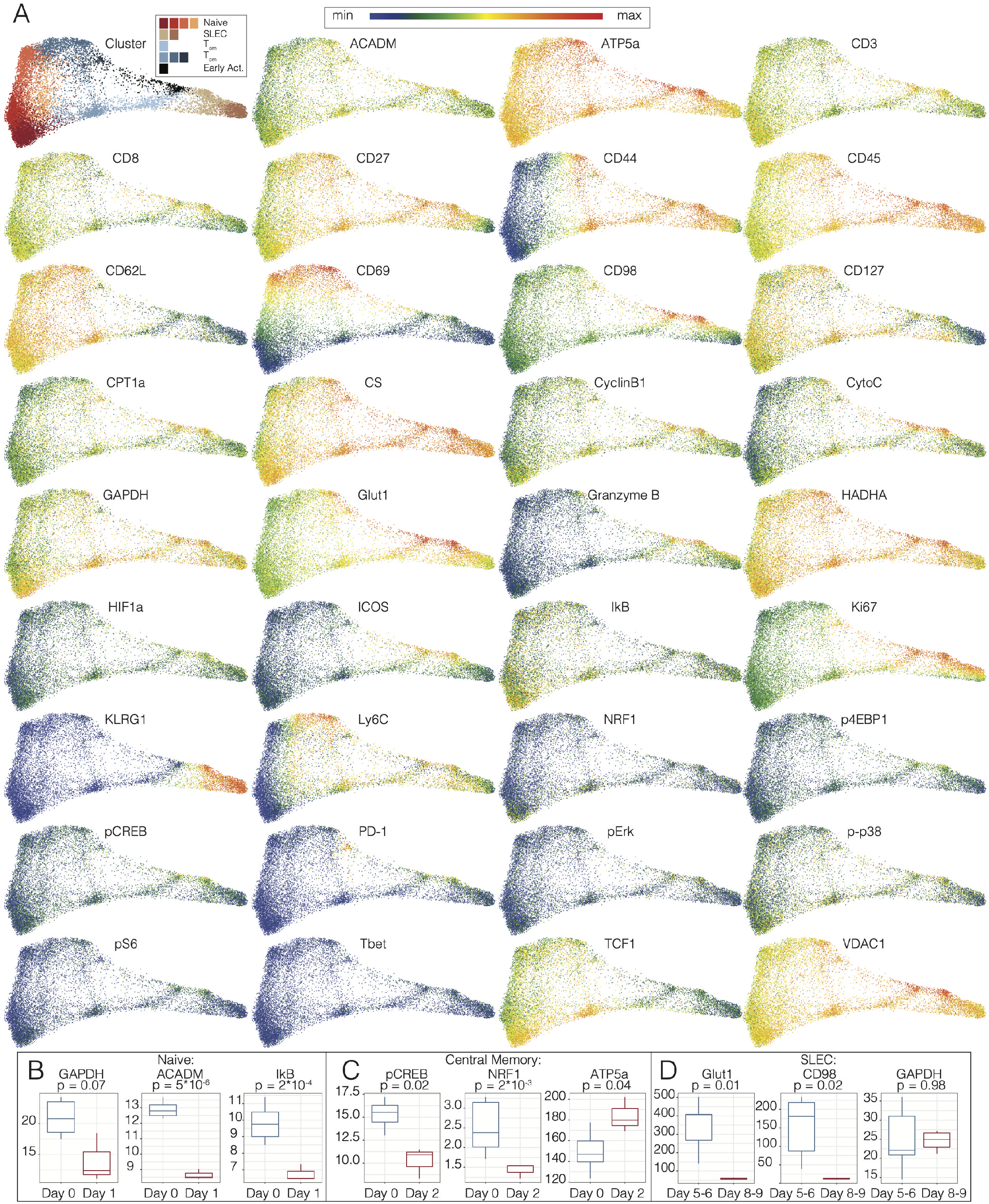
Single-cell metabolic analysis of the primary CD8 T cell response in vivo. **(A)** Pooled CD8 T cells from mice at days 0 to 9 of Lm-OVA infection (n=2-3 mice per time point) clustered by Phenograph and visualized by force-directed graphs. Force-directed graphs of single-cell expression profiles of individual markers are depicted. Box plots of marker expression of **(B)** naïve, **(C)** central memory, and **(D)** SLECs at specified time points post-infection with Listeria monocytogenes. Significance analysis by paired two-tailed student’s t-test. Whiskers represent 1.5 * IQR.

**Figure S3.**
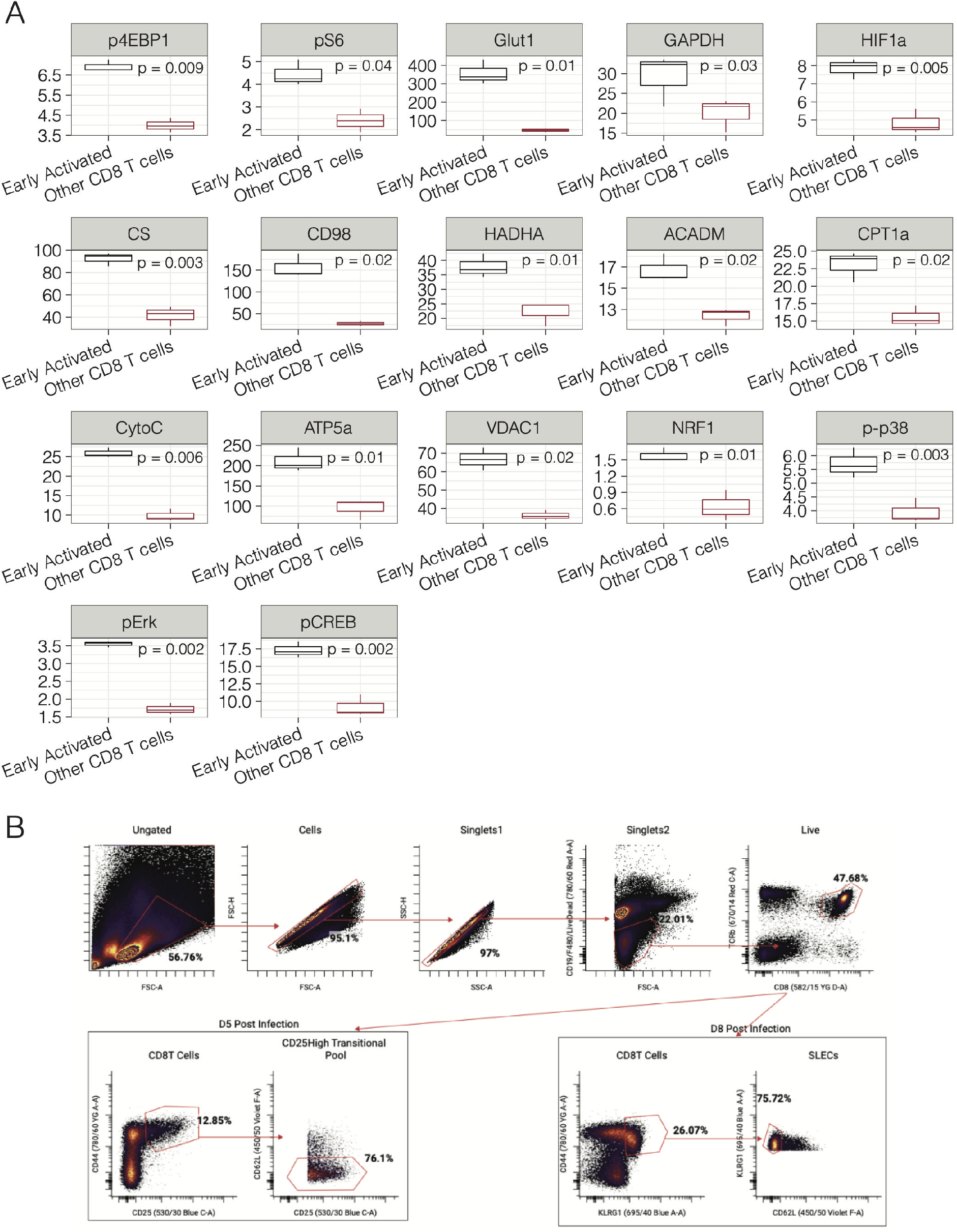
Single-cell metabolic analysis by mass cytometry reveals the unique metabolic profile of early activated CD8 T cells. **(A)** Box plots of marker expression of early activated T cells compared to all other CD8 T cells at day 5 p.i. with Listeria monocytogenes. Significance analysis by paired two-tailed student’s t-test. Whiskers represent 1.5 * IQR. (B) Sorting strategy for isolation of naïve (CD62L^hi^CD44^low^KLRG1^low^CD25^lo^), transitional effectors (CD62L^low^CD44^hi^CD25^hi^) at day 5 post-infection and SLECs (CD62L^low^CD44^hi^KLRG1^hi^CD25^low^) and day 8 post-infection is depicted by biaxial plots.

**Figure S4.**
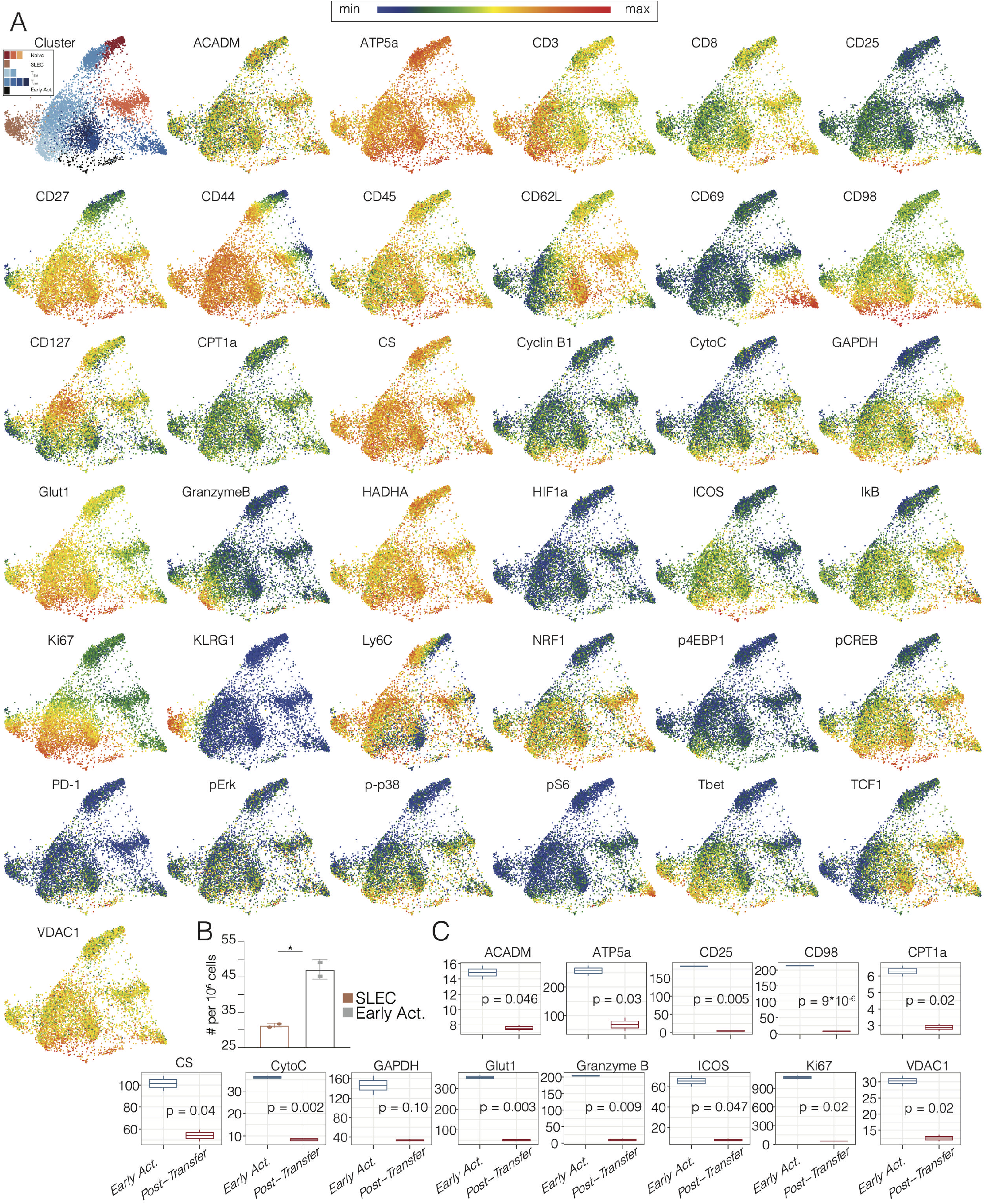
Single-cell metabolic analysis of the antigen-specific primary CD8 T cell response in vivo. **(A)** Pooled OT1 CD8 T cells from mice at days 3 to 7 of Lm-OVA infection (n=1-3 mice per time point) clustered by Phenograph and visualized by a force-directed graphs. Force-directed graphs of single-cell expression profiles of individual markers are depicted. **(B)** Number of congenic transferred cells derived from either early activated cells or SLECs four days after adoptive transfer. **(C)** Metabolic and signaling marker profiles of OT1 CD8 T cells before and after transfer at days 5 and 9 p.i. respectively are represented by histograms. Significance analysis by paired two-tailed student’s t-test. Whiskers represent 1.5 * IQR.

